# Histology-aware Inference of Cell-cell Communication in Spatial Transcriptomics

**DOI:** 10.64898/2026.01.22.701166

**Authors:** Xiaofei Wang, Chenyang Tao, Yu Jiang, Yixuan Jiang, Hanyu Liu, Zheng Jiang, Pinan Zhu, Ningfeng Que, Jianzhong Xi, Stephen Price, Binzhi Qian, Jianguo Xu, Chao Li

## Abstract

Spatial transcriptomics (ST) enables inference of cell-cell communication (CCC) from ligand-receptor expression and spatial proximity, but these measurements incompletely capture the histological context that may facilitate or restrict signaling. Here we introduce HARMONIC, a histology-aware graph learning framework integrating ST and paired hematoxylin and eosin (H&E)-stained images to contextualize candidate CCC using cellular morphology and tissue architecture. HARMONIC quantifies conditional statistical dependencies among molecular, cellular morphological and tissue architectural features, and integrates these representations through self-supervised graph learning. In synthetic benchmarks, HARMONIC recovers interactions supported by permissive tissue contexts despite weak molecular evidence and suppresses spurious predictions across architectural barriers. Across diverse tissues and ST platforms, HARMONIC identifies anatomically concordant communication patterns at tumor-immune interfaces, around necrosis, across renal compartments and within tumor-associated microvasculature, with complementary transcriptional and spatial proteomic support. HARMONIC demonstrates how histological context can refine spatial CCC inference beyond molecular compatibility and proximity.

## Introduction

Cell-cell communication (CCC) underpins tissue organization and function across development, homeostasis and disease, often through ligand-receptor (LR) interactions between sender and receiver cells^1^. Spatially resolved CCC inference typically uses spatial transcriptomics (ST) to infer candidate LR interactions from cellular transcriptomic profiles and spatial proximity^2–8^. Yet these molecular and spatial measurements alone do not fully capture the tissue context in which signaling occurs. Paired hematoxylin and eosin (H&E) images provide complementary information on cellular morphology and tissue architecture, but these histological cues remain underutilized as the explicit contextual information for CCC inference^3–6,9–14^.

At the cellular level, H&E-derived morphology provides phenotypic information complementary to transcriptomic profiles in characterizing potential sender and receiver cells^15^. For instance, mitotic figures can indicate proliferative activity, a cellular process influenced by intercellular signaling^16^. Integrating cellular morphology with gene expression may therefore provide a more informative representation of candidate interacting cells.

Beyond individual cells, ST provides limited direct information about tissue architecture, which can constrain or facilitate intercellular signaling^6,17^. For instance, anatomical features such as compartment boundaries (e.g., basement membranes) and luminal spaces (e.g., blood vessel interiors) can physically separate cells that are close in spatial coordinates, making some proximity-based interactions less anatomically plausible^18,19^. Conversely, extracellular scaffolds can retain signaling molecules within local niches and support localized communication^20,21^. H&E-based characterization of tissue architecture can therefore provide complementary anatomical context for assessing the biological plausibility of inferred interactions.

Here we introduce HARMONIC, a histology-aware graph learning framework that integrates ST and paired H&E images to infer LR-mediated CCC within tissue context. HARMONIC incorporates cellular morphology and surrounding tissue architecture as contextual information for sender-receiver interactions, alongside spatial expression data. It first organizes neighboring cells with similar transcriptomic and morphological characteristics into local niches, then uses conditional mutual information (CMI) to quantify statistical dependencies among cellular gene expression and H&E-derived cellular morphological and tissue architectural features at the niche level. These context-aware representations are subsequently integrated using a self-supervised graph learning approach, resulting in refined CCC scores.

We benchmarked HARMONIC against representative spatial CCC inference approaches^4–6,22^, using simulations and real-world datasets. In simulations, HARMONIC recovered interactions with weak molecular evidence in permissive tissue contexts while suppressing spurious predictions across architectural barriers. We further evaluated HARMONIC across distinct tissue settings: (i) cellular interfaces, where in ovarian cancer, H&E images delineated tumor-immune boundaries, allowing HARMONIC to infer anatomically aligned immune-tumor signaling; (ii) structural transitions, where around necrotic regions, H&E images resolved necrotic cores and tumor rims under-represented by ST, enabling the model to distinguish spatial patterns of migration- and proliferation-associated signaling; (iii) anatomical compartments, where in mouse kidney, H&E-defined renal zones and glomeruli, facilitating recovery of anatomically compartmentalized signaling, especially in regions with sparse LR co-detection; and (iv) microvascular structures, where in a newly acquired papillary craniopharyngioma (PCP) cohort, H&E-resolved microvasculature allowed more precise localization of tumor-endothelial signaling compared to ST-only methods.

Together, HARMONIC demonstrates how incorporating histological context can refine spatial CCC inference beyond molecular compatibility and proximity, enabling more anatomically informed investigations of intercellular interactions in complex tissues.

## Results

### 2.1 Model framework, synthetic and robustness validation

#### 2.1.1 HARMONIC overview

HARMONIC is built on a cell-cell graph *G* = (**V**, **E**), where each node *v* ∈ **V** encodes cellular features and each edge *e* ∈ **E** represents the CCC score between a sender-receiver cell pair (Fig. 1a-c). Because cellular morphology alone may not distinguish different cell types or states, node features integrate transcriptomic profiles from ST with cellular morphological features from H&E images, extracted using both a segmentation-based morphology extractor adapted from MorphLink^15^ (with features defined in Table S1) and a pathology foundation model^24^. Edge representations were constructed by feature-wise concatenation of cellular expression and morphological features, intercellular distance, and H&E-derived tissue architectural features at local and global scales (see *Methods*). These representations are optimized through the histology-aware graph learning process described below (Fig. 1d-f).

**Fig. 1:**
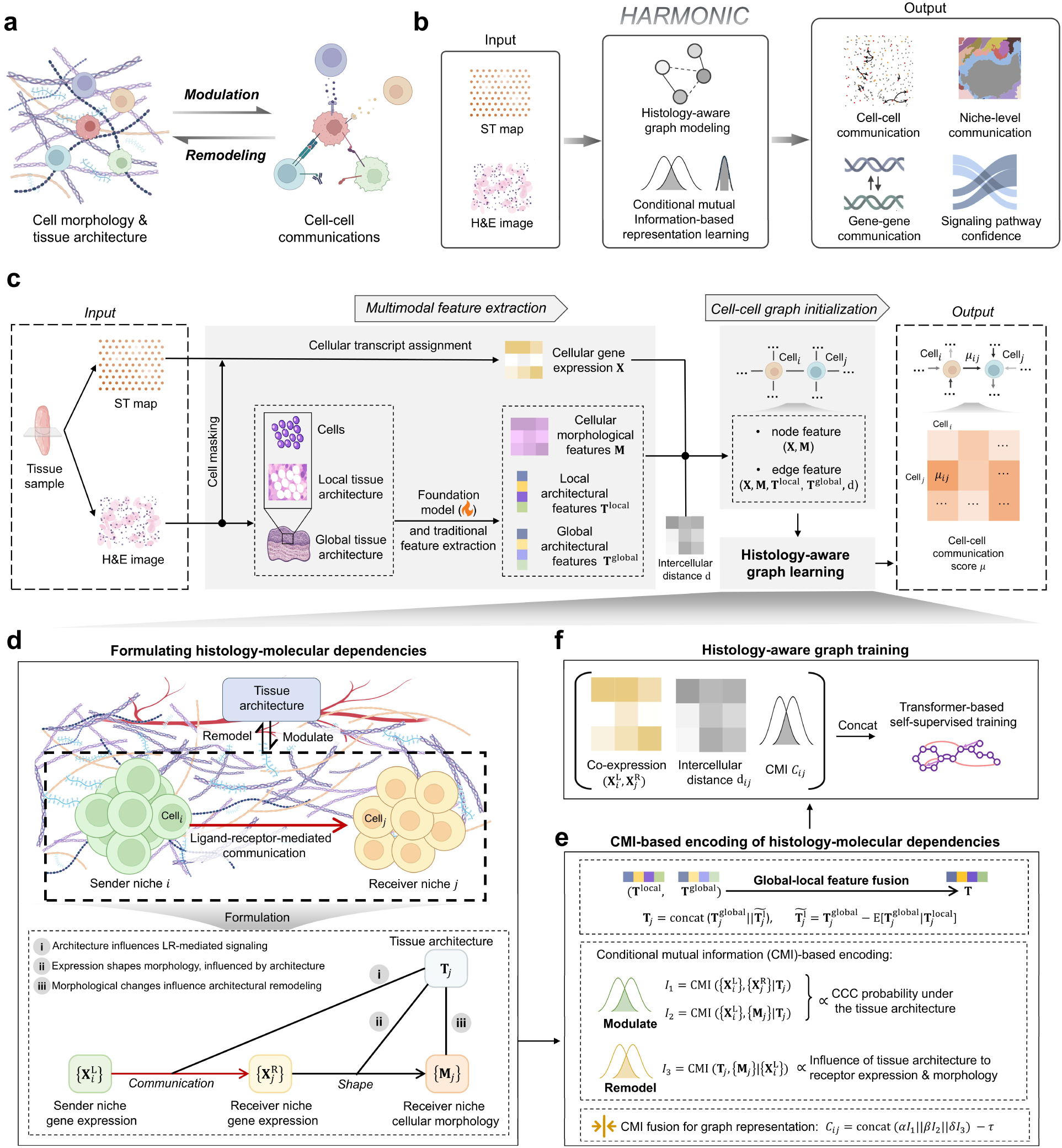
Study design and model overview. **a**, **Conceptual illustration** of the relationships among cell-cell communication, cellular morphology and tissue architecture. **b**, **Study design**. HARMONIC takes the multimodal input of paired ST and H&E images, with the output of spatially resolved CCC. **c**, **HARMONIC framework.** Based on the H&E-derived cell mask, HARMONIC extracts multimodal features from ST and H&E images, used for constructing the cell-cell graph *G* = (**V**, **E**), with nodes **V** indicating cellular features and edges **E** denoting CCC scores. Edge features are further optimized via a histology-aware graph learning process to obtain final CCC scores. **d-f**, **Histology-aware graph learning process. d**, For each sender-receiver pair, a histology-aware graph (HAG) modulates the communication score at the niche level by capturing biological dependencies among tissue architecture, cellular gene expression and cell morphology. **e**, Conditional mutual information is used to formulate these dependencies within the HAG, which are incorporated through a transformer-based attention mechanism to refine the communication score. **f**, Finally, we optimize all parameters via a masked-edge reconstruction^23^-based self-supervised learning strategy tailored to CCC inference.

**First**, HARMONIC considers three complementary, biologically motivated relationships between molecular activity and tissue organization (Fig. 1d): **(i)** tissue architecture can influence LR-mediated signaling, in part by shaping cell contacts and ligand accessibility^20^; **(ii)** sender-cell ligand expression may be associated with receiver-cell morphology, potentially reflecting signaling-related cellular programs influenced by local tissue geometry and mechanics^25^; **(iii)** changes in cell morphology and collective cellular organization, including cell shape and packing, can contribute to tissue architectural remodeling^26^. We model these relationships at the niche level to reduce sensitivity to low signal-to-noise ratios in ST and cell segmentation errors in H&E images, which can compromise cellular transcript assignment^5,27^ and morphological representation^28^. Hence, multimodal spatial clustering^17^ identifies niches comprising spatially adjacent cells with similar transcriptomic and morphological features.

**Next**, HARMONIC formalizes these relationships as conditional statistical dependencies using conditional mutual information (CMI) (Fig. 1e), a measure widely used in graphical representation learning^29^ and biological network inference^30^. Specifically, each dependency is formulated by a dedicated CMI term; for example, the conditional dependence on tissue architecture of communication from a sender niche *i* to a neighboring receiver niche *j* under a given LR pair is defined as:

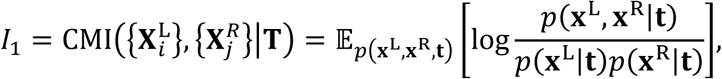

where 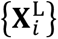 and 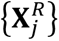 denote ligand and receptor expressions in sender and receiver niches, respectively, while **T** represents the tissue architectural features derived by fusing nested global (**T**^global^) and local (**T**^local^) architectural representations. This term models the dependence between ligand and receptor expression conditional on the surrounding tissue architecture, allowing co-expression evidence to be evaluated within its histological context. The remaining two dependencies are captured by their respective CMI terms, as detailed in *Methods*. The three CMI terms are weighted by learnable parameters and concatenated to form a fused CMI representation. For each candidate sender-receiver cell pair, HARMONIC identifies the niches containing the two cells and incorporates the corresponding niche-level CMI features into its edge representation.

**Finally**, HARMONIC refines these CMI-based edge representations by aggregating information from neighboring edges through a transformer-based multi-head cross-attention module^31^. The refined representation is used to compute a CCC score for the corresponding LR pair and sender-receiver cell pair. Parameters are optimized with a masked-edge reconstruction^23^-based self-supervised objective tailored to CCC inference. The contribution of each dependency is further assessed through ablation analyses in Section 2.1.3.

**Extended Fig. 1:**
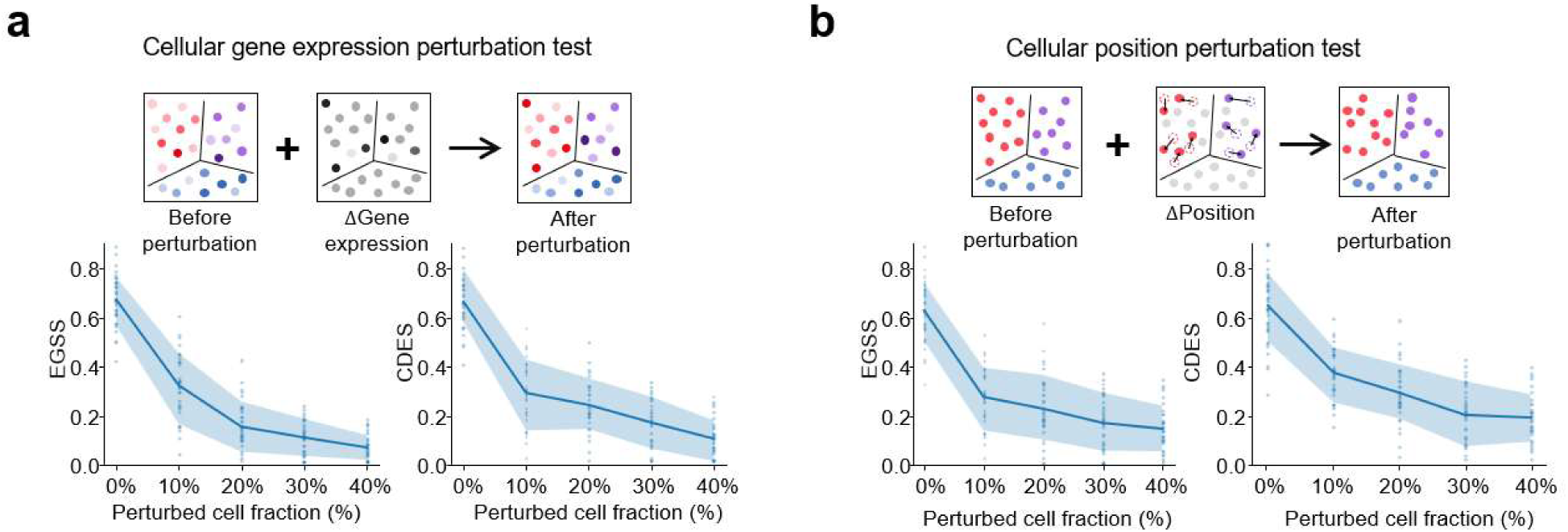
Sensitivity of inferred CCC to molecular and spatial perturbations in mouse brain. **a-b,** Cellular gene-expression perturbation (**a**) and cell-position perturbation (**b**) tests, evaluating with increasing fractions of perturbed cells. Both metrics progressively decreased with stronger perturbation, confirming that the inferred CCC patterns depend on coherent cellular molecular profiles and spatial organization. Results are averaged across cells and LR pairs from CellChatDB group^34^.

#### 2.1.2 Synthetic validation of HARMONIC in rectifying ST-only CCC inference

To assess the contribution of histological context in CCC inference under controlled conditions, we generated synthetic data representing two complementary scenarios: (**I**) recovering underestimated communication between senders and receivers with low LR co-expression due to the low ST signal-to-noise ratio; and (**II**) suppressing spurious communication across physical barriers, such as tissue-compartment boundaries or vascular lumens. Of note, in both scenarios, the expression of each receiver cell is designed to be jointly determined by neighboring senders and the surrounding tissue architecture (*Methods*).

**Scenario I** modeled tissue architecture associated with enhanced CCC, analogous to settings such as the interface of tumor and immune infiltration where tumor-immune interactions are often enriched. We compared HARMONIC with its *ST-only* variant, which does not model tissue architecture, to determine whether architectural context enables recovery of otherwise missed interactions. For sender-receiver pairs with relatively high LR co-expression, both methods detected ground-truth interactions. However, when LR co-expression was simulated at reduced levels (fivefold lower; *Methods*), the ST-only variant of HARMONIC missed substantially more ground-truth interactions (Fig. 2a). HARMONIC recovered these interactions, reducing the mean false-negative rate from 0.820 (95% CI 0.787-0.853) to 0.271 (0.252-0.289; Fig. 2b). This improvement persisted across a continuous range of architecture-mediated promotion strengths (Fig. S1), supporting robustness to different architectural effects. A real-world validation example is shown in Section 2.5.2.

**Fig. 2:**
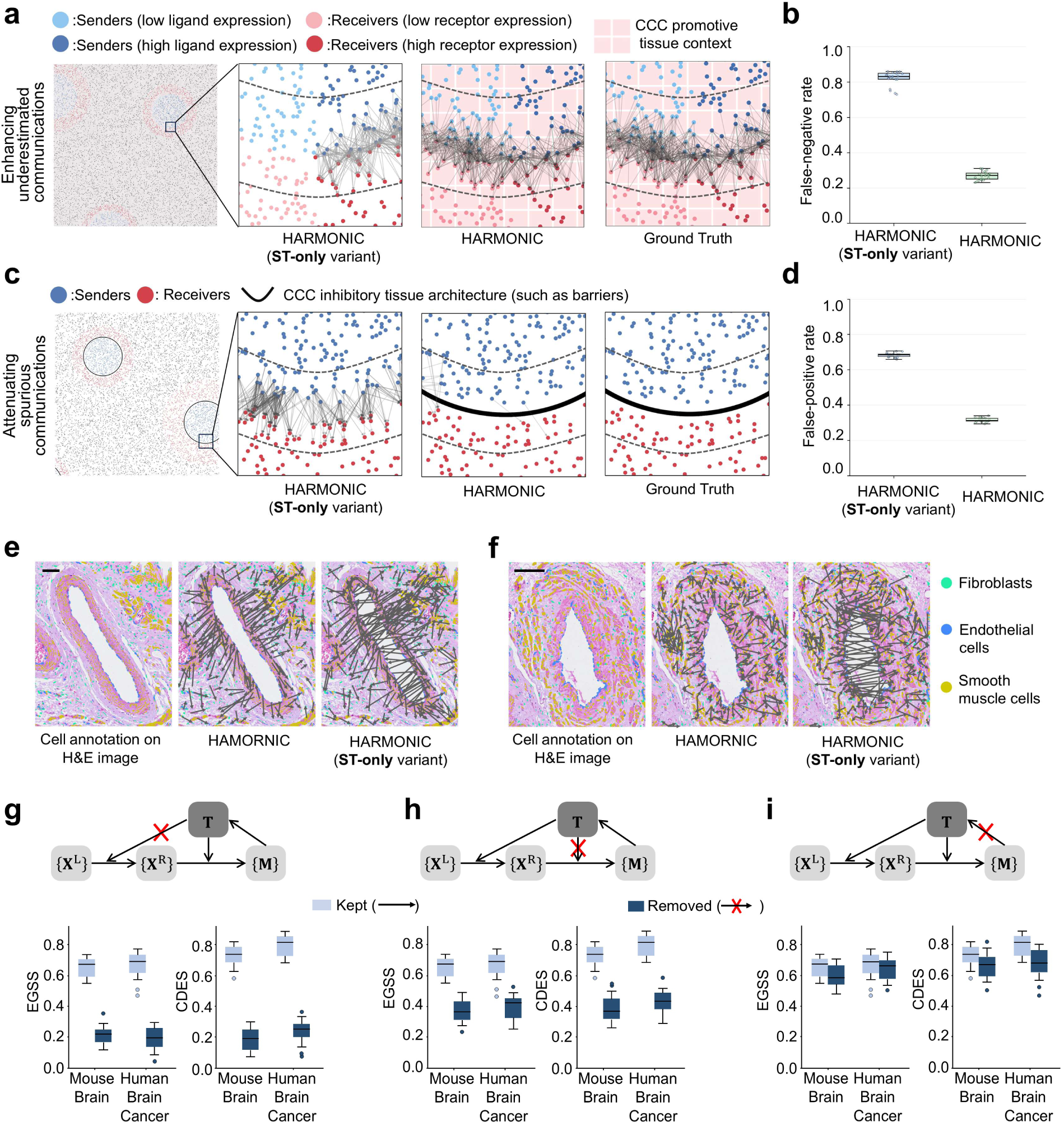
Effects of histological context on simulated CCC inference and sensitivity analyses. **a-d**, **Synthetic validation**. **a,** CCC facilitated by tissue context. In a synthetic setting where permissive tissue context supports communication between sender and receiver cells with relatively weak LR co-expression, HARMONIC recovers context-supported interactions underestimated by its ST-only variant, more closely matching the ground truth. Dashed lines indicate the maximum sender-receiver communication distance. See *Methods* for simulation details. **b,** False-negative rates of CCC inference for HARMONIC and its ST-only variant. **c,** Communication constrained by tissue context. When tissue context restricts communication between neighboring sender and receiver cells, the ST-only variant of HARMONIC infers spurious interactions, whereas HARMONIC suppresses them, more closely matching the ground truth. **d,** False-positive rate for HARMONIC and its ST-only variant. **e-f, Real-world examples.** Representative vascular regions showing inferred CCC among fibroblasts, endothelial cells and smooth muscle cells. Compared with its ST-only variant, HARMONIC reduces cross-luminal interactions and better conforms to vascular architecture. Scale bar: 100µm. **g-i**, **Robustness analysis**. Dependency-ablation analysis of the histology-molecular dependencies used by HARMONIC, evaluated on EGSS^32^ and CDES^33^. Ablating individual dependencies generally reduces both metrics, supporting their contributions to robust CCC inference.

**Scenario II** simulated an architectural barrier that could restrict cross-barrier CCC. The *ST-only* variant predicted numerous false-positive interactions across the barrier, probably as ST alone does not encode this structural constraint (Fig. 2c). By explicitly incorporating tissue architecture, HARMONIC suppressed most spurious predictions and reduced the false-positive rate from 0.683 (95% CI, 0.678-0.689) to 0.317 (0.311-0.322; Fig. 2d). This improvement was maintained across varying strengths of architecture-mediated signal blocking (Fig. S2). We next tested whether this finding extended to a biologically realistic architectural barrier by examining vascular structures (Fig. 2e-f). We focused on *PDGFB-PDGFRB*, a canonical endothelial-perivascular stromal signaling axis^35^. The arterial lumen within a stromal region of human ovarian cancer tissue is approximately 100 μm in diameter, creating an anatomical separation expected to constrain short-range and contact-dependent signaling across the lumen^36,37^. The *ST-only* variant produced many cross-lumen interactions, whereas HARMONIC largely reduced them.

Together, these controlled experiments indicate HARMONIC’s potential in improving CCC inference fidelity over ST-only methods, with explicit incorporation of histological context.

#### 2.1.3 Dependency ablation and perturbation sensitivity analyses

To investigate the contribution of the three tissue-architecture-related biological dependencies in HARMONIC, we ablated each dependency individually while retaining the other two and compared CCC inference before and after ablation. Evaluations were performed across healthy (mouse brain and kidney) and disease (human brain and ovarian cancer) tissues (Fig. 2g-i; Fig. S3). To assess the molecular and histological concordance of inferred CCC, we utilized the contextual distance enrichment score^33^ (CDES; *Methods*). We also assessed inference stability using the edge-level graph stability score^32^ (EGSS; *Methods*), which quantifies the preservation of inferred CCC under transcriptomic perturbations.

Ablation reduced both metrics across all the tested tissues, with the largest decrease occurring after removal of the tissue-architecture dependency that modulates LR-mediated signaling. Specifically, EGSS decreased by 0.44, 0.51, 0.49 and 0.37, whereas CDES decreased by 0.55, 0.37, 0.56 and 0.48 in mouse brain, mouse kidney, human brain cancer and ovarian cancer, respectively (Fig. 2g; Fig. S3a).

We next assessed the sensitivity of HARMONIC predictions to controlled molecular and spatial perturbations by independently introducing Gaussian noise into cellular gene-expression profiles and spatial coordinates, following previous perturbation-based analyses^38^. As the fraction of perturbed cells increased from 0% to 40% in mouse brain, EGSS and CDES decreased from 0.68 and 0.64 to 0.07 and 0.11, respectively, under gene-expression perturbation, and to 0.16 and 0.19 under spatial-coordinate perturbation (Extended Fig. 1). Similar trends were observed in mouse kidney and ovarian cancer (Fig. S4). These results characterize the sensitivity of the inferred CCC networks to increasing disruption of molecular and spatial information.

**Extended Fig. 2:**
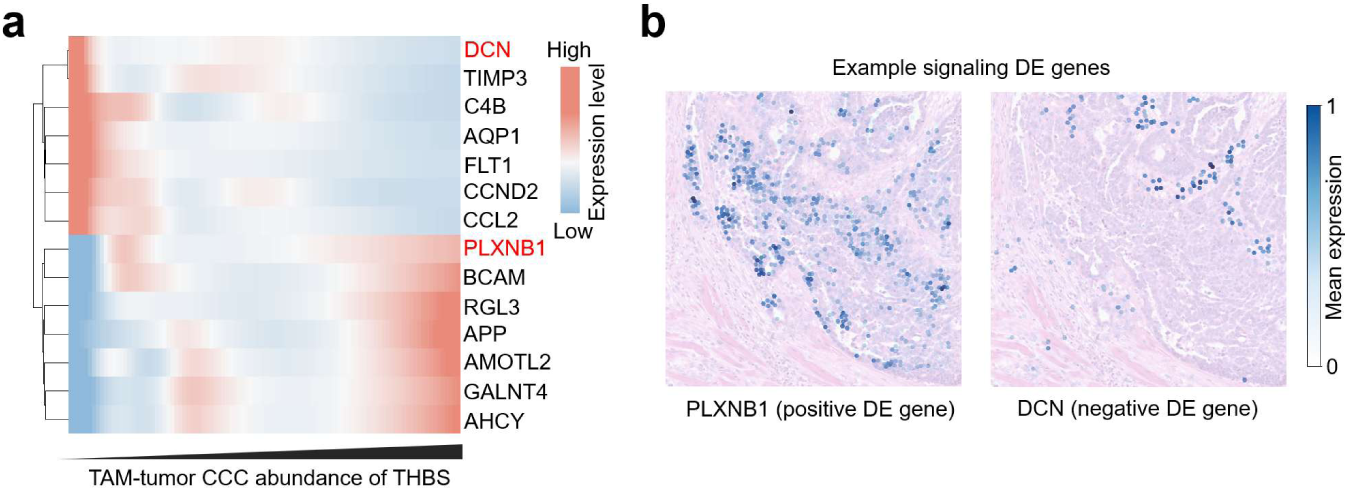
Biological concordance of HARMONIC-inferred THBS signaling. **a,** Differentially expressed (DE) genes associated with HARMONIC-detected TAM-tumor THBS signaling. **b**, Spatial maps of representative positive and negative DE genes.

### 2.2 HARMONIC resolves directional TAM-tumor communication at the interface of tumor and immune infiltration

Immune-infiltrated tumor regions contain densely intermingled tumor and immune cells, where close cellular packing can complicate cell delineation and cellular transcript assignment, potentially confounding ST-only CCC inference^27^. H&E provides complementary cellular morphological and tissue architectural information, enabling delineation of the physical interface of tumor and immune infiltration (Fig. 3a). We therefore applied HARMONIC to an ovarian cancer Xenium sample^42^ to enhance the biological fidelity of inferred tumor-immune CCC.

**Fig. 3:**
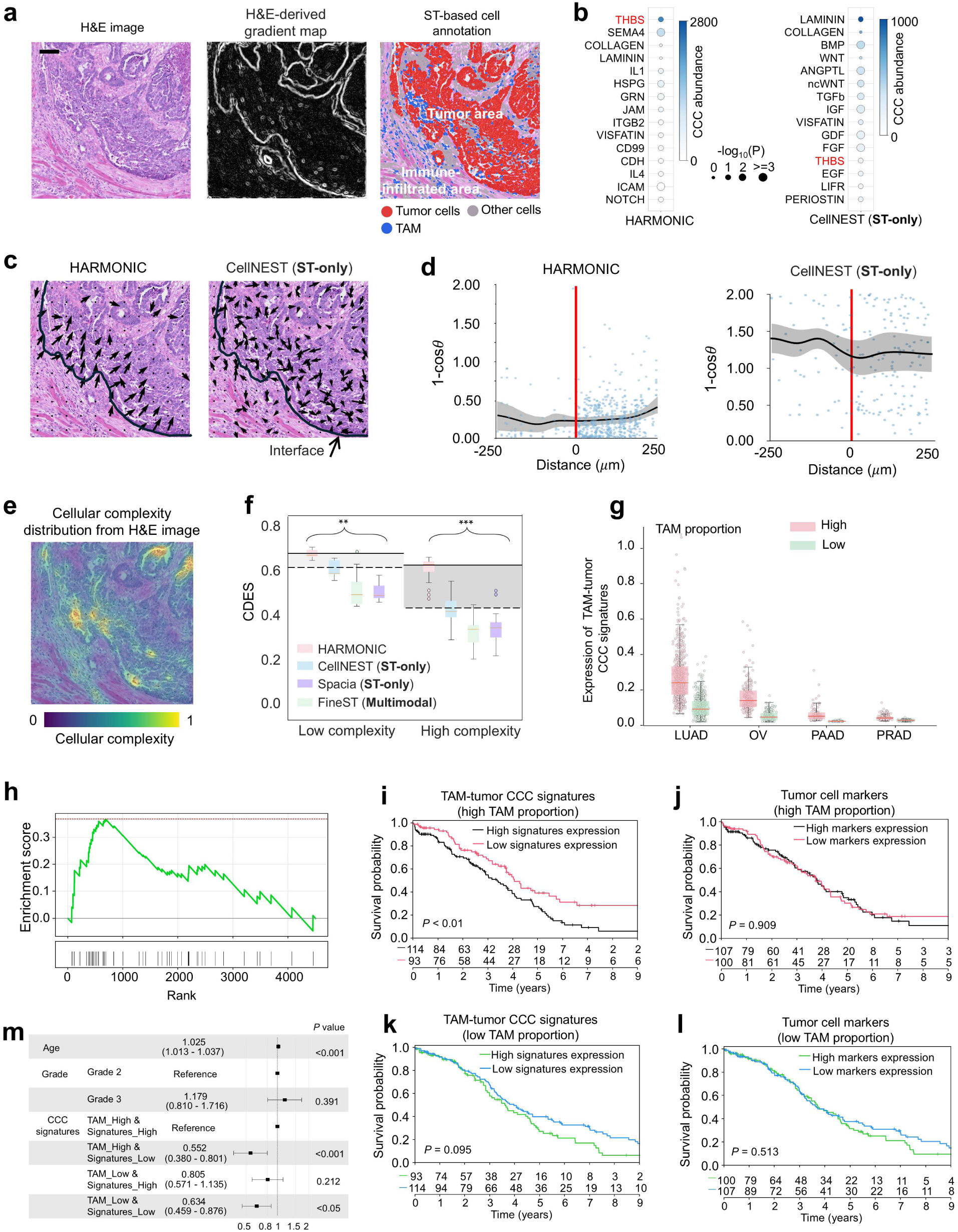
HARMONIC reveals directional TAM-tumor CCC at the interface of tumor and immune infiltration. **a,** H&E image and its gradient map showing tissue architecture, alongside ST-based annotations of cell distributions. **b,** CellChatDB^34^ pathways ranked by CCC abundance from HARMONIC and ST-only baseline, CellNEST^6^, with THBS most abundant in HARMONIC, consistent with its reported role in TAM-associated tumor adhesion and progression^39^. **c-d,** Spatially inferred THBS communication directions (**c**) and their alignment with the interface (**d**), quantified by 1 − cos *θ*, where lower values indicate stronger alignment. HARMONIC showed stronger directional alignment than CellNEST. **e,** H&E-derived cellular complexity^40^, showing higher complexity at the interface. **f,** CCC inference performance in low- and high-complexity regions, evaluated by CDES; HARMONIC showed higher CDES than tested ST-only and multimodal methods, particularly in highly complex regions. **g, E**xpression of HARMONIC-predicted TAM-tumor signature in the TCGA samples, stratified by TAM proportions (LUAD, N=518; OV, N=434; PAAD, N=183; PRAD, N=554), showing consistently higher receptor expression in TAM-high tumors. **h,** GSEA in TCGA-OV comparing tumors with high and low TAM proportions, showing enrichment of the inferred TAM-tumor signature in TAM-high tumors. **i-l,** Prognostic value of the HARMONIC-predicted TAM-tumor signature (**i**, TAM-high; **k**, TAM-low) compared with established tumor-cell markers^41^ (**j**, TAM-high; **l**, TAM-low) in TCGA-OV, showing stronger overall-survival stratification by the TAM-tumor signature, particularly in patients with high TAM proportions. **m,** Multivariable Cox models adjusting for age and TCGA-reported cancer grade, with TAM-tumor signature remaining significant in TAM-high patients. Scale bar: 100µm.

Based on ST-derived cell annotation, tumor cells (n=4,172) and tumor-associated macrophages (TAMs; n=737) were identified as the predominant cell populations at the interface of tumor and immune infiltration (Fig. S5a). Given their abundance and close spatial association (Fig. S5a; Fig. 3a), we examined TAM-tumor cell communication across this interface. As shown in Fig. 3b and Fig. S5b, HARMONIC identified THBS as the most abundant signaling pathway, consistent with previous evidence linking THBS signaling to TAM-associated tumor adhesion and progression^39^. By contrast, the representative **ST-only** method CellNEST^6^ detected substantially weaker THBS signaling (Fig. 3b).

Then, to assess the transcriptional patterns associated with HARMONIC-inferred THBS signaling, we used tradeSeq^4,43^ to identify tumor cell genes whose expression varied with received THBS signaling, treating THBS abundance as a continuous covariate analogous to pseudotime-based differential expression analysis^44^ (Extended Fig. 2). *PLXNB1* showed the strongest positive association with THBS signaling, whereas *DCN* showed the strongest negative association (Wald test; BH-adjusted *P*=1.9×10^-171^ for *PLXNB1* and 1.3×10^-85^ for *DCN*; Extended Fig. 2a). Spatially, increased *PLXNB1* expression corresponded to stronger THBS signaling, whereas *DCN* expression showed the opposite pattern (Extended Fig. 2b; Fig. S6). These patterns are consistent with previous reports associating high *PLXNB1* and low *DCN* expression with tumor adhesion and progression^44,45^. Applying the same analysis to other abundant HARMONIC-inferred pathways, including SEMA4 and COLLAGEN, revealed similarly concordant tumor cell expression patterns (Figs. S7-S8), further supporting the transcriptional concordance of the inferred CCC.

We next examined whether incorporating H&E-derived tissue architecture could improve the spatial organization of inferred CCC. HARMONIC inferred directionally coherent THBS signaling across the interface of tumor and immune infiltration (Fig. 3c) that aligned with H&E-derived morphological gradients (Fig. 3a). These signals closely followed boundary-normal directions from TAMs toward tumor cells, with a mean angular difference of 33° compared with 118° for the **ST-only** baseline CellNEST, where 0° indicates perfect alignment and 180° the opposite direction (paired Wilcoxon signed-rank test, *P*<10^-5^; Fig. 3d).

Further, we examined whether incorporating tissue architecture provided greater benefit in architecturally complex regions. The interface showed higher tissue complexity than the remaining regions (complexity score^40^: 0.724 versus 0.346; Fig. 3e). Although HARMONIC performed comparably to ST-only and multimodal baselines^5,6,22^ in low-complexity areas, HARMONIC showed clear advantages in high-complexity regions across CCC metrics of CDES^33^ and EGSS^32^ (HARMONIC 0.64/0.58 versus CellNEST 0.43/0.53, Spacia 0.36/0.51, FineST 0.35/0.46; all *P*<0.001; Fig. 3f; Fig. S9), supporting the value of histological context for CCC inference in complex tissue environments.

Finally, we evaluated the cross-cancer robustness of HARMONIC-identified TAM-tumor signatures (Table S2), defined as receptor genes in the top five TAM-tumor cell signaling pathways, including THBS, COLLAGEN, BMP, LAMININ and IL1. Specifically, we assessed whether the TAM-tumor signatures showed concordant patterns across cancer types using The Cancer Genome Atlas (TCGA) bulk RNA-seq data from lung, ovarian, pancreatic, and prostate cancers. BayesPrism^46^ estimated TAM proportions and tumor-cell-specific gene expression of TAM-tumor signatures for each patient. Across all four cancer types, higher TAM proportions were associated with higher tumor-cell receptor expression (Fig. 3g; two-sided Wilcoxon rank-sum test, *P* (lung cancer) = 5.05×10^-131^, *P* (ovarian cancer) = 1.11×10^-59^, *P* (pancreatic cancer) = 1.25×10^-30^, *P* (prostate cancer) = 4.49×10^-79^), providing cross-cancer support for the HARMONIC-identified TAM-tumor THBS axis.

### 2.3 HARMONIC-derived TAM-tumor signatures are associated with patient survival

We further assessed the prognostic relevance of HARMONIC-predicted TAM-tumor signatures. We first examined whether these signatures captured TAM-associated transcriptional states in tumor cells using RNA-seq data from TCGA ovarian cancer samples (n=414). We deconvolved^46^ the RNA-seq data to cell-type-specific gene expression, and compared differential expression in the tumor-cell component between patients with high and low TAM proportions. Gene-set enrichment analysis (GSEA)^47^ suggested that TAM-tumor signatures identified by HARMONIC were significantly enriched in the top differentially expressed genes (*P*=0.04; Fig. 3h), supporting their association with tumor-cell states linked to TAM-tumor signaling.

We next performed Kaplan-Meier analyses based on HARMONIC-predicted TAM-tumor signatures in tumor cells, with established tumor-cell markers^41^ (Table S3) as the control (Fig. 3i-l). Because the signatures were derived from TAM-tumor communication, we hypothesized that their prognostic association would be more evident in tumors with higher TAM infiltration. We therefore stratified patients into TAM-high and TAM-low groups based on BayesPrism-estimated TAM proportions. We found that high expression of the TAM-tumor CCC signature indeed predicted worse survival in the TAM-high patients (*P*<0.01), whereas there were no significant associations for the TAM-low patients (*P*=0.095). By contrast, the expression level of tumor-cell markers^41^ was less prognostic in both patient groups (TAM-high: *P* =0.909; TAM-low: *P*=0.513). In multivariable Cox models adjusting for age and TCGA-reported cancer grade, the TAM-tumor signatures remained significantly associated with overall survival in TAM-high patients (*P*<0.001; hazard ratio=0.552, 95%CI=0.380-0.801, with high signature expression as the reference; Fig. 3m; Table S4). These findings support the prognostic value of HARMONIC-derived TAM-tumor CCC signatures.

### 2.4 HARMONIC characterizes spatially organized tumor migration and proliferation activities around necrotic areas

Necrotic regions are hallmarks of solid tumors and can influence surrounding tumor-cell activities, including migration, invasion and proliferation^49^. However, CCC inference around necrosis remains challenging from ST alone, as molecular expression and cellular proximity in ST maps do not explicitly capture necrotic boundaries and the associated tissue organization^50^. H&E images delineate necrotic cores, viable tumor rims and surrounding tissue architecture (Fig. 4a), providing ST-complementary structural context for reliable CCC inference. We therefore applied HARMONIC to paired Xenium ST and H&E data from an ovarian cancer sample (4,305 cells; 5,101 genes) to characterize spatially organized CCC around necrotic regions.

**Fig. 4:**
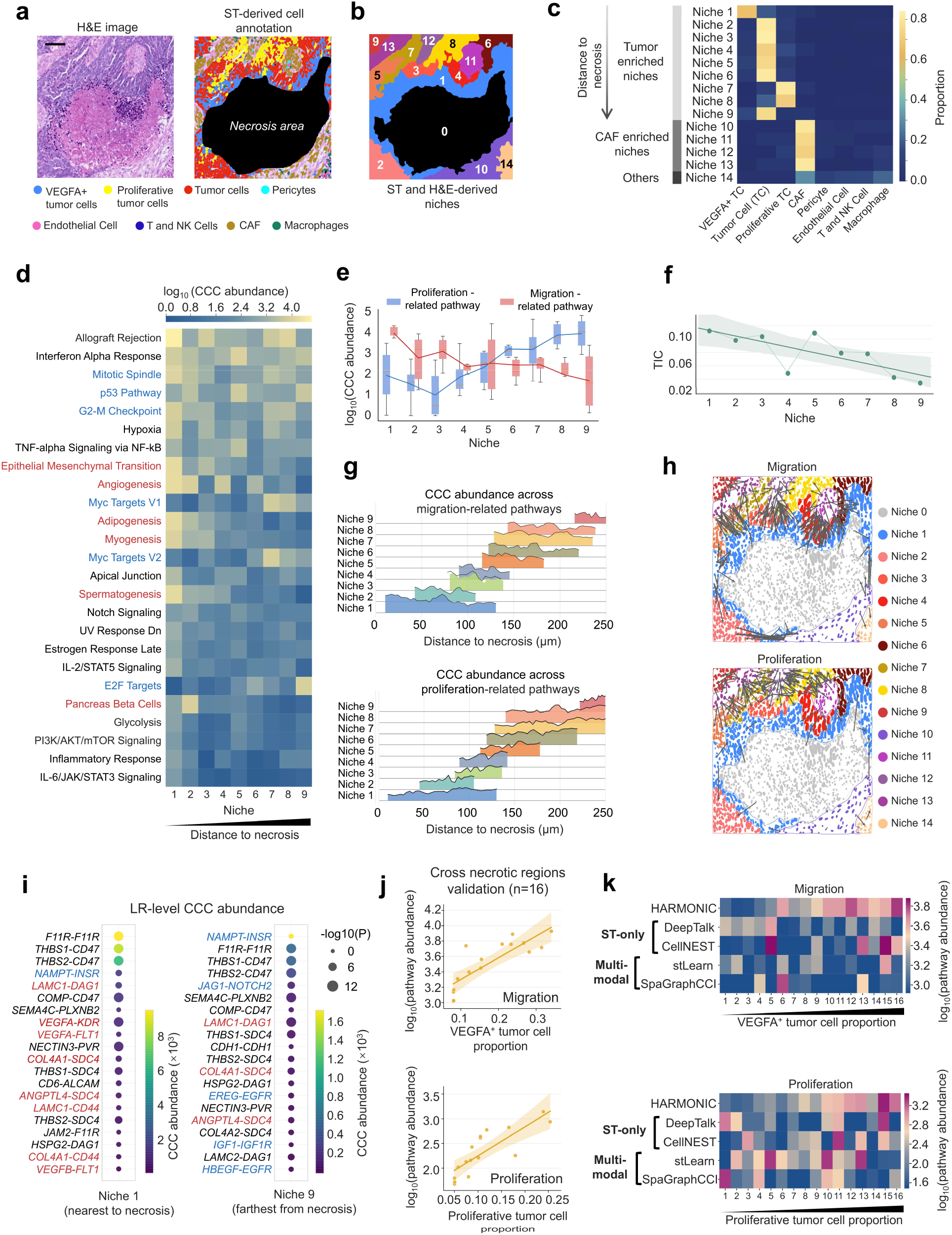
HARMONIC reveals spatially organized tumor activities around necrotic areas. **a,** H&E image and ST-derived cell annotation of a representative necrotic area in human ovarian cancer. **b,** Spatial niches defined from multimodal ST-H&E features. **c,** Cell-type composition across niches; niches 1-9, ordered by increasing distance from necrosis, are tumor-enriched, whereas niches 10-13 are CAF-enriched. **d,** CCC abundance of the top 50% Hallmark MSigDB^48^ pathways across tumor-enriched niches. Migration- and proliferation-related pathways are highlighted in red and blue, respectively. **e-f**, Overall CCC abundance of migration- and proliferation-related pathways (**e**) and tumor-initiating cell (TIC) proportion (**f**) across niches, revealing stronger migration-related signaling and higher TIC abundance near necrosis, with proliferation-related signaling increasing with distance. **g-h,** Within-niche distribution (**g**) and spatial localization (**h**) of migration- and proliferation-related CCC along the distance-to-necrosis gradient. **i**, LR-level CCC abundance in niches 1 (nearest to necrosis) and 9 (farthest from necrosis), with migration-related LR pairs (red) more enriched in niche 1 and proliferation-related pairs (blue) in niche 9. **j-k**, Validation across all necrotic regions (n=16) in the same sample. HARMONIC-derived migration- and proliferation-related CCC abundance increases with the proportions of VEGFA^+^ and proliferative tumor cells, respectively (**j**), whereas this correspondence is less evident for ST-only and multimodal methods (**k**). Scale bar: 100µm.

HARMONIC identified 14 spatially organized cellular niches, with most niches enriched in tumor cells (n=10) or cancer-associated fibroblasts (CAF; n=4) (Fig. 4b-c). Given the prominence of these two compartments, we next examined whether CAF-tumor communication varied across the necrosis-associated tissue gradient, along which tumor-cell states are known to change markedly^49^. Tumor-enriched niches closer to necrosis showed greater CAF-tumor communication involving migration- and angiogenesis-associated pathways^48^, including epithelial-mesenchymal transition and angiogenesis (Fig. 4d; Fig. S10a). Consistently, the proportions of tumor-initiating cells and *VEGFA*^+^ tumor cells decreased with distance from necrosis, showing concordant spatial variation with migration- and angiogenesis-associated CCC (Fig. 4a, f). In contrast, proliferation-associated pathways, including E2F Targets and G2-M Checkpoint programs, showed stronger abundance in more distal niches (Fig. 4d; Fig. S10a), accompanied by an increasing proportion of *MKI67*^+^ proliferative tumor cells (Fig. S10b). Within-niche signaling patterns showed the same spatial trends (Fig. 4g), with representative CCC networks illustrated in Fig. 4h. Together, these results reveal a spatial transition from migration- and angiogenesis-associated signaling near necrosis toward proliferation-associated signaling with increasing distance from the necrotic core.

Beyond pathway-level analysis, we examined LR-level CCC abundance in niches. The niche closest to necrosis (niche 1) showed greater representation of migration- and angiogenesis-associated interactions, including *ANGPTL4-SDC4 and VEGFA-KDR*. By contrast, the most distal tumor-enriched niche (niche 9) showed greater representation of proliferation-associated interactions, including *EREG-EGFR* (Fig. 4i). These LR-level differences were consistent with the broader spatial variation in Hallmark-associated CCC programs and supported the biological interpretability of the inferred interactions.

To assess the spatial consistency of these patterns, we extended the analyses to all 16 necrotic regions within the same ovarian cancer sample (cells= 29,840; genes= 5,101). Across regions, the abundances of migration- and proliferation-associated CCC programs inferred by HARMONIC correlated with the proportions of *VEGFA*^+^ and proliferative tumor cells, respectively (Fig. 4j). By contrast, **ST-only methods**, including DeepTalk^22^, CellNEST^6^, as well as multimodal baselines, including stLearn^19^ and SpaGraphCCI^13^, showed weaker concordance between CCC and cell states (Fig. 4k). These results indicate greater concordance between HARMONIC-inferred CCC programs and local tumor-cell states across spatially distinct regions.

### 2.5 HARMONIC reveals anatomically compartmentalized signaling in mouse kidney

#### 2.5.1 Zone-specific signaling across major renal regions

To evaluate HARMONIC in a highly compartmentalized tissue, we applied it to a Visium HD adult mouse kidney sample (cells=277,728, genes=18,993). The tissue comprised four transcriptionally distinct regions, i.e., inner medulla (IM), inner/outer stripe of the outer medulla (ISOM/OSOM) and cortex (Fig. 5a). This anatomical organization is clearly captured in H&E images (Fig. 5b), providing complementary structural context for region-associated CCC inference.

**Fig. 5:**
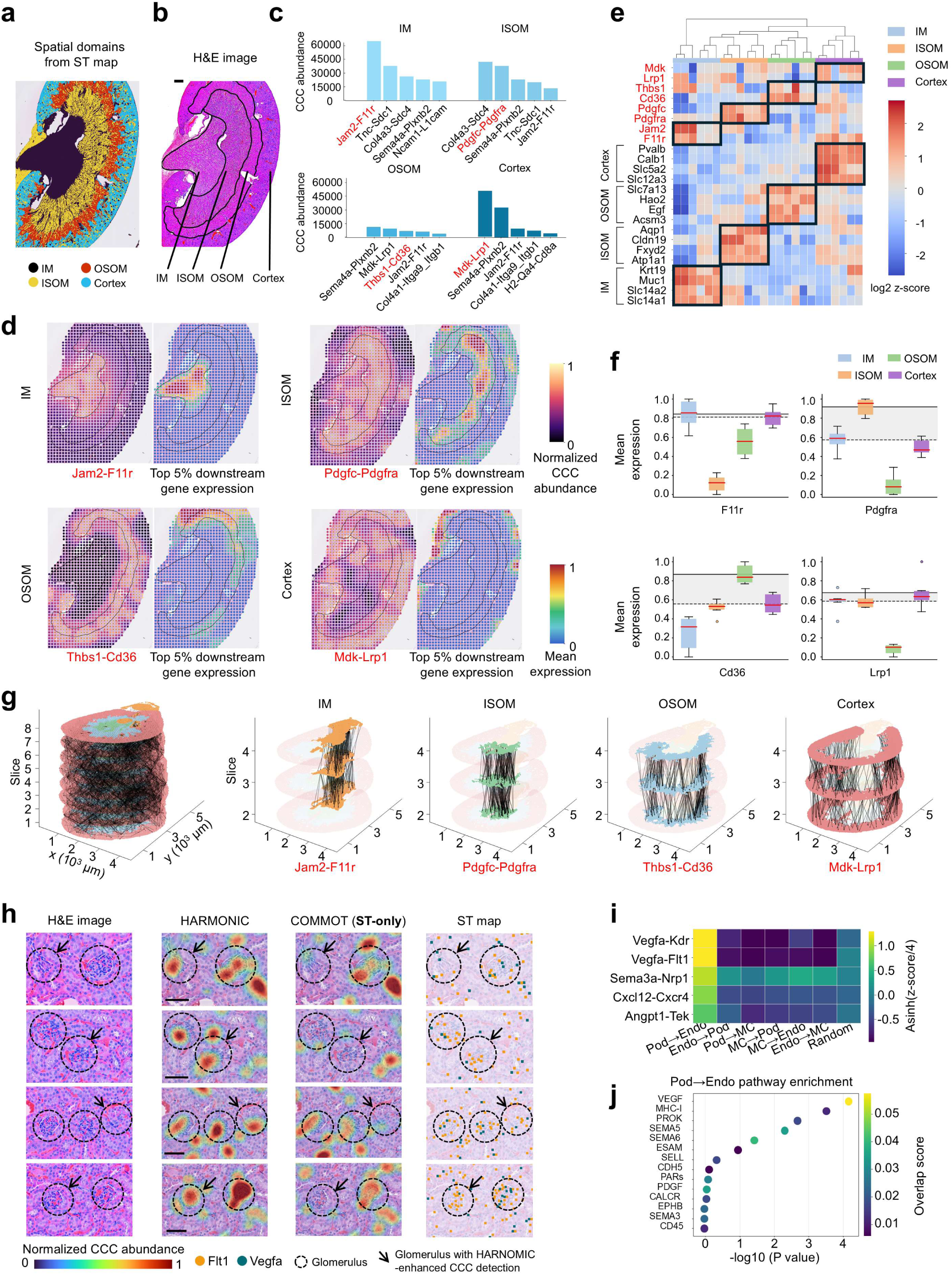
HARMONIC recovers signals in regions with distinct morphology in mouse kidney. **a-b**, ST-derived spatial domains (**a**) and corresponding H&E-defined renal zones (**b**), including cortex, OSOM, ISOM and IM. **c**, Most abundant inferred CCC within each zone, highlighting *Jam2-F11r* in IM, *Pdgfc-Pdgfra* in ISOM, *Thbs1-Cd36* in OSOM and *Mdk-Lrp1* in cortex. **d**, Spatial distributions of these interactions and expression of their top downstream genes, showing concordant zone-restricted patterns. **e**, Heatmap displaying LR gene expression and canonical regional markers (Table S5), averaged across six ST samples. **f**, Box plots showing mean expression (n=6) levels of receptors in detected LR pairs in distinct regions, supporting the anatomical specificity of the inferred signaling. **g**, 3D reconstruction across serial sections showing preservation of zone-specific CCC patterns across tissue depth, supporting the applicability of HARMONIC to emerging 3D ST. **h-j**, **Recovering CCC inference in glomeruli**. **h**, Representative glomeruli comparing *Vegfa-Flt1* communication inferred by HARMONIC and ST-only baseline, COMMOT, with the underlying ST map. HARMONIC more closely localizes signaling to glomerular structures, especially in glomeruli with weaker LR co-expression in ST. **i,** Directional glomerular CCC across major cell-type pairs, showing preferential podocyte-to-endothelial VEGF signaling. **j,** Pathway enrichment of podocyte-to-endothelial communication, with VEGF showing the strongest enrichment. Scale bar: 100µm. MC: mesangial cell. Pod: podocyte. Endo: endothelial cell.

We first assessed whether HARMONIC could identify reliable region-associated LR pairs. In the cortex, HARMONIC reported 193 LR pairs out of 2,017 total pairs in the top 20% strongest signals, with *Mdk-Lrp1* being the most abundant (Fig. 5c), primarily between proximal and distal convoluted tubule epithelial cells. Notably, *Mdk* encodes a heparin-binding growth factor in renal proximal tubules in the cortex, whereas *Lrp1* is an established receptor for *Mdk*^51^. In the ISOM, *Pdgfc-Pdgfra* was found abundant, which was established to promote renal interstitial fibroblast proliferation and fibrosis^52^. Other representative kidney-associated pairs included *Thbs1-Cd36* in the OSOM, a proapoptotic matricellular axis implicated in ischemic proximal-tubule injury^53^, and *Jam2-F11r* in the IM, whose components regulate epithelial/endothelial junction integrity and leukocyte transmigration^54^. To assess transcriptional concordance with these inferred LR interactions, we mapped the mean expression of the top 5% of predicted downstream genes associated with each representative LR pair. These gene-expression patterns broadly colocalized with the corresponding CCC signals across renal regions (Pearson’s correlation coefficient [PCC]=0.67, *P*<0.001, Fig. 5d), providing complementary transcriptional support for the inferred spatial patterns.

To further assess the regional transcriptional support for HARMONIC-inferred LR signals, we analyzed six independent mouse kidney ST samples from 10X Genomics and aggregated spatial expression within each anatomical region to generate regional pseudobulk profiles (Fig. 5e-f). Across samples, *Lrp1* expression was significantly higher in the cortex than in IM, ISOM, and OSOM (Friedman test with Holm-adjusted Wilcoxon post hoc tests, *P*=0.013; Fig. 5e-f). Likewise, *F11r*, *Pdgfra*, and *Cd36* were enriched in the IM, ISOM, and OSOM, respectively (IM, *P*=2.19×10^-3^; ISOM, *P*=1.03×10^-3^; OSOM, *P*=4.26×10^-3^; Fig. 5e-f), providing independent transcriptional evidence for the corresponding regional LR signals. **By contrast, ST-only** methods, e.g., Spacia and COMMOT, showed less consistent recovery of region-associated CCC (Figs. S11-12), indicating HARMONIC’s ability to identify region-associated CCC.

Finally, we examined the consistency of these region-associated CCC patterns across tissue depth using an independent 3D Array-seq mouse kidney dataset^55^ comprising eight serial sections (92,769 spots; 15,144 genes). HARMONIC identified *Mdk-Lrp1*, *Thbs1-Cd36*, *Pdgfc-Pdgfra*, and *Jam2-F11r* in the cortex, OSOM, ISOM and IM, respectively, across consecutive sections (Fig. 5g; Fig. S13). The recurrence of region-associated LR patterns across eight serial sections supports their spatial consistency along tissue depth and illustrates the extension of HARMONIC to serial-section spatial transcriptomic data.

#### 2.5.2 Glomerulus-specific signaling within renal microanatomy

In mouse kidney, glomerular homeostasis involves interactions between podocytes and glomerular endothelial cells, including *VEGF*-family signaling. However, limited local detection of ligand and receptor transcripts in individual glomeruli may weaken ST-only inference of candidate interactions. We therefore investigated whether incorporating histological context could improve the anatomical localization of podocyte-to-endothelial communication, focusing on the *Vegfa-Flt1* axis.

By incorporating H&E-derived glomerular boundaries and surrounding renal architecture, HARMONIC identified localized candidate *Vegfa-Flt1* interactions from podocytes to endothelial cells within morphologically delineated glomeruli, including regions with relatively weak but detectable LR evidence (Fig. 5h). In these regions, the **ST-only** method COMMOT generally produced weaker inferred communication scores. These findings indicate that histological context can help prioritize more anatomically plausible interactions when local molecular evidence is limited.

Beyond *Vegfa-Flt1*, we also observed other canonical LR pairs within glomeruli, including *Vegfa-Kdr*, a key regulator of glomerular endothelial survival and fenestration^56^, and *Cxcl12-Cxcr4*, an established signal for glomerular vascular development^57^. Most signals occurred between podocytes and endothelial cells (Fig. 5i). Pathway enrichment analysis of genes involved in abundant LR pairs in podocyte-endothelial signaling identified VEGF as the most significantly enriched pathway, together with vascular and intercellular signaling programs including SEMA5/6 and ESAM (Fig. 5j). These findings suggest the biological relevance of HARMONIC-inferred CCC within glomerular microanatomy.

### 2.6 HARMONIC identifies tumor-endothelial communication in rare tumor of papillary craniopharyngioma

PCP is a rare benign suprasellar tumor with a highly vascularized microenvironment^58^. Its dense microvascular architecture can be delineated by H&E images, providing structural context for reliable CCC inference. To evaluate whether HARMONIC can resolve biologically meaningful communication in this rare tumor, we first analyzed an index Xenium PCP sample from our cohort (cells=29,095, genes=5,001; Fig. 6a). Among 1,939 inferred LR pairs, the ten most abundant tumor-endothelial pairs included several angiogenic and vascular-remodeling signals, notably *VEGFA-KDR*, the most abundant, *JAG1-NOTCH1* and *EFNB2-EPHB4* (Fig. 6b). *VEGFA-KDR* communication scores were spatially concentrated around vascular structures (Fig. 6c), consistent with a perivascular distribution of inferred tumor-endothelial interactions. These scores were also positively associated with the expression of predicted downstream genes (Fig. 6d; *Methods*), providing transcriptional concordance with the inferred *VEGFA-KDR* signaling pattern.

**Fig. 6:**
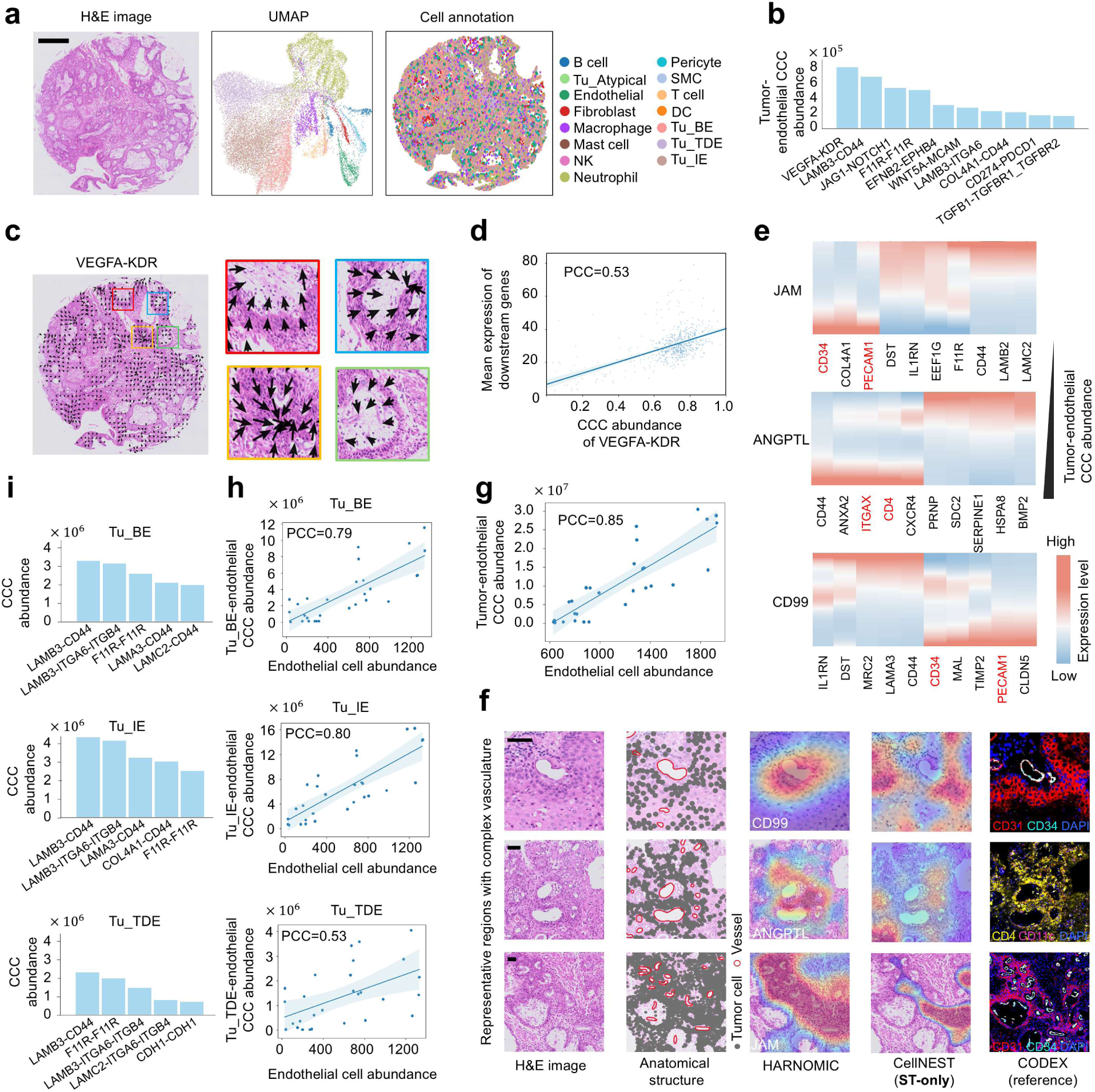
HARMONIC identifies tumor-endothelial communication in PCP. **a**, H&E image, UMAP, and ST-derived cell annotation. **b**, Top 10 LR pairs ranked by inferred tumor-endothelial CCC abundance, with *VEGFA-KDR* showing the highest abundance. **c**, Spatial vector field^4^ of inferred *VEGFA-KDR* signals, with zoomed views of four representative vessel-rich regions, supporting a spatially coordinated VEGF signaling program. **d**, Correlation between *VEGFA-KDR* CCC abundance and mean downstream-gene expression, with each dot representing a cell (PCC=0.53). **e**, Heatmaps of top 10 differentially expressed (DE) genes of 3 pathways (JAM, ANGPTL, and CD99) with known important roles in tumor-endothelial interactions. **f**, Representative regions with complex vasculature comparing HARMONIC and CellNEST against anatomical and CODEX references. HARMONIC more closely localized tumor-endothelial signaling to vascular structures. **g**, Correlation between endothelial cell abundance and tumor-endothelial CCC abundance across 29 PCP samples; each dot represents one PCP sample (PCC=0.85). **h-i**, Correlation between endothelial cell abundance and tumor-endothelial CCC abundance (**h**) and top 5 abundant LR pairs (**i**) across three tumor subtypes, revealing state-specific tumor-endothelial signaling programs. Scale bar: 100µm.

Next, at the pathway-level, we examined three pathways with known important roles in tumor-endothelial interactions, namely the JAM^59^, ANGPTL^60^, and CD99^61^ pathways. We identified genes whose expression varied with increasing received signaling and modeled their expression trends (Fig. 6e). Increasing JAM signaling was associated with higher expression of the vascular endothelial markers *CD34* and *PECAM1* and lower expression of the basement-membrane-associated genes *LAMB2* and *LAMC2*, consistent with JAM’s known role in vascular barrier regulation^59^. CD99 showed a similar pattern, whereas increasing ANGPTL signaling was associated with higher *ITGAX* and *CD4* expression, suggesting an immune-enriched state in ANGPTL-active perivascular niches. In representative regions with complex vasculature, HARMONIC localized these tumor-endothelial signals more closely to vessel-associated structures than the **ST-only** CellNEST, with greater spatial concordance with vessel anatomy and CODEX staining (Fig. 6f).

Moreover, to assess the robustness of these observations across the cohort, we extended the analysis to all 29 Xenium PCP samples from 19 patients (cells=935,881, genes=5,001; Fig. S14). Across samples, total tumor-endothelial CCC abundance strongly correlated with endothelial cell abundance (PCC=0.85, *P*=5.02×10^-9^; Fig. 6g), supporting the biological consistency of the inferred tumor-endothelial communication.

Finally, we investigated tumor-endothelial communication across the four transcriptionally defined tumor states that we recently identified^62^: basal-like epithelial state (Tu_BE, PCC=0.89, *P*=4.03×10^-7^), intermediate epithelial state (Tu_IE, PCC=0.79, *P*=1.53×10^-7^), terminally differentiated epithelial state (Tu_TDE, PCC=0.53, *P*=3.25×10^-3^; Fig. 6h), and atypical epithelial state (Tu_Atypical, PCC=0.68, *P*=4.47×10^-5^; Fig. S15). Subtype-specific LR rankings revealed partially shared but quantitatively distinct CCC programs: tumor-vascular adhesion-related *LAMB3-CD44* and *LAMB3-ITGA6* signaling recurred across subtypes, whereas the remaining leading LR pairs varied by subtype (Fig. 6i).

## Discussion

Inferring intercellular communication using LR pairs is important for deciphering cellular activity in tissue. While existing scRNA-seq-based methods for identifying CCC in tissue samples cannot resolve communication between individual cells due to the lack of cell-cell proximity information, recent advances in single-cell-resolved ST technologies now enable CCC inference at the individual cell level. However, ST alone incompletely represents non-cellular tissue architecture and provides limited information on cellular morphology. As a result, ST-only approaches may infer interactions that are transcriptomically plausible but less anatomically plausible, such as predictions across physical barriers like basement membranes or vascular lumina.

Recent CCC inference methods, including CellNEST^6^ and GITIII^10^, have increasingly modeled the local neighborhoods of sender-receiver cell pairs, but derive this context primarily from ST-derived molecular profiles and spatial coordinates, without explicitly modeling histological tissue architecture. In addition, GITIII^10^ infers sender-to-receiver influences on receiver-cell transcriptional states rather than resolving communication at the level of LR pairs. In parallel, multimodal ST-histology frameworks have increasingly integrated transcriptomic and histological information to improve spatial resolution, cellular representation and tissue characterization^11,12,14,19^. For example, FineST^11^ uses H&E-derived features to enhance the ST spatial resolution and subsequently identifies LR interactions from the enhanced transcriptomic representation, rather than directly incorporating histological architecture into CCC inference.

Despite these advances, tissue architecture remains incompletely modeled in spatial CCC inference as the explicit contextual constraint of whether candidate sender-receiver interactions are supported or constrained by their anatomical environment. HARMONIC addresses this gap by directly incorporating cellular morphology and surrounding tissue architecture into the scoring of candidate LR-mediated interactions, enabling molecular and spatial evidence to be interpreted within their histological context. HARMONIC uses conditional mutual information to model dependencies among tissue architecture, LR-mediated signaling, cellular gene expression and morphology, and integrates these context-aware representations through graph learning.

Controlled simulations highlighted the complementary roles of histological context in CCC inference: supporting candidate interactions with limited molecular evidence yet permissive tissue context, and constraining proximity-based predictions across anatomical barriers. These findings suggest that histological context does not simply strengthen or weaken inferred communication, but can influence how molecular and spatial evidence is interpreted according to the surrounding tissue environment. Thus, similar LR expression and spatial proximity may support different interaction predictions depending on local tissue organization.

Applications across tissues further show how architectural context can refine the interpretation of spatial signaling and help recover biologically plausible CCC otherwise missed by ST-only methods. At the interface of tumor and immune infiltration, inferred signaling directions aligned with histological boundaries, suggesting that these interfaces act as active communication zones. Around necrotic regions, HARMONIC revealed a spatial transition from migration- and angiogenesis-related signaling near necrosis to proliferative signaling in more distal areas. Similarly, the recovery of anatomy-specific signaling in mouse kidney and tumor-endothelial interactions in PCP indicate that architectural context can reveal CCC patterns that are difficult to resolve from transcript abundance alone. Together, these findings show how explicitly incorporating cellular morphology and tissue architecture can refine the anatomical interpretation of candidate LR-mediated interactions.

HARMONIC provides a flexible framework for incorporating complementary histological information into CCC inference. However, several limitations remain. First, anatomical concordance supports the biological plausibility of predicted interactions but does not establish functional LR signaling. Targeted genetic perturbation of implicated ligand or receptor genes, coupled with downstream signaling assays, could provide more direct functional validation. In addition, CMI captures conditional statistical associations rather than causal relationships. Future extensions could explore causal modeling and incorporate additional relationships involving receptor activation, downstream signaling and multicellular feedback. Moreover, more efficient and interpretable H&E representations could prioritize signaling-relevant tissue structures while reducing redundant gigapixel-scale information. Finally, the framework could also extend beyond predefined databases to propose context-specific or previously unannotated LR pairs for experimental validation.

## Methods

### 1. HARMONIC framework

#### Overview

HARMONIC is a histology-aware graph learning framework for inferring directed LR-mediated CCC using paired ST and H&E images. The inputs comprise a cell-by-gene expression matrix, cell coordinates, a spatially registered H&E image, cell masks and an LR database. For each LR pair, HARMONIC constructs a directed cell-level graph in which edges represent candidate sender–receiver interactions selected by spatial proximity and LR activity, rather than established signaling events. The model then integrates cell-level molecular and morphological features with H&E-derived tissue architecture and niche-level conditional mutual information (CMI) representations through an edge-biased graph Transformer^63^. Following masked-edge self-supervised training, HARMONIC assigns an LR-specific CCC score to each candidate edge for ranking and downstream analysis.

#### Data preprocessing and histology-aware feature extraction

For each sample, the processed cell-by-gene expression matrix was obtained after ST quality control and normalization. The paired H&E image was registered to the ST coordinate system and segmented to obtain cell masks, thereby linking each cell *i* to its expression profile **x**_i_, spatial coordinate **s**_i_, and corresponding histological region.

HARMONIC extracted histological features at two complementary levels. Cellular morphology **m**_i_ combined interpretable descriptors derived from the cell mask with appearance embeddings extracted from a cell-centered H&E patch using a pretrained pathology image encoder. Tissue architecture was represented by nested, receiver-centered H&E contexts at local and global scales. By default, these contexts covered 256 *μ*m and 1,280 *μ*m, respectively, or the maximum available field of view when the tissue was smaller. This setting is consistent with prior studies^22,64^. The local representation was obtained from a target-cell-removed patch nested within the global context, reducing direct reuse of the target cell’s appearance as architectural evidence. The local and global embeddings, denoted 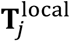 and 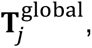, were retained separately and fused to form the receiver-centered tissue-architecture representation **T**_j_. Feature-extraction procedures and hyperparameters are provided in Supplementary section 1.

#### LR-specific candidate communication graph construction

For each LR pair (L, R), HARMONIC constructed a directed graph *G*^(L,R)^ = (**V**, **E**^(L,R)^), where each node represents an individual cell and each edge connects a candidate sender-receiver pair. A directed edge from cells *i* and cell *j* is included when the cells satisfy the predefined spatial-neighborhood criterion and ligand *L* and receptor *R* meet their activity criteria in the sender and receiver cells, respectively. These edges define the candidate interactions to be evaluated, rather than confirmed signaling events. Each candidate edge is initially represented by the intercellular distance *d_ij_*, LR co-expression strength 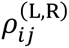 and an LR-identity embedding **q**^(L,R)^. H&E-derived tissue architectural features and niche-level CMI representations are subsequently incorporated into these edge features during histology-aware graph learning (Step 3).

#### Histology-aware graph learning

HARMONIC linked niche-level estimation of histology-molecular dependencies to cell-level graph scoring. Cellular niches provided robust organizational units for estimating CMI, whereas the graph Transformer retained individual cells as nodes to generate cell-resolved, directed CCC inferences.

##### *Step1*: Niche construction

Cellular niches were delineated by multimodal spatial clustering of the paired ST and H&E data. For each cell, the normalized transcriptomic profile, H&E-derived morphological features, and spatial coordinates were concatenated into a joint representation and clustered using *k*-means. To suppress isolated labels that were inconsistent with the surrounding tissue, cells within a 60-*μ*m radius were treated as a spatial neighborhood and cluster labels were smoothed by local majority voting. This procedure produced spatially contiguous niches composed of neighboring cells with similar transcriptomic and morphological states. The resulting niches were used to estimate the histology-molecular dependencies described below.

##### *Step2:* CMI-based formulation of histology-molecular dependencies

Consider a candidate communication from sender niche N_i_ to an adjacent receiver niche N_j_ under LR pair (L, R). Let 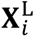 and 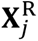 denote ligand and receptor expression in the sender and receiver niches, respectively; **X**_j_ the receiver-niche transcriptomic state; **M**_j_ the receiver-niche morphological representation; and **T**_j_ the fused local-global tissue-architecture representation around the receiver niche. For random variables *a*, *b* and *c*, CMI is defined as:

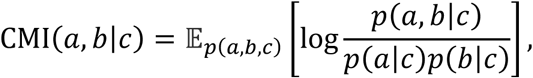

Under a multivariate Gaussian approximation, CMI can be estimated from the corresponding conditional covariance matrices as:

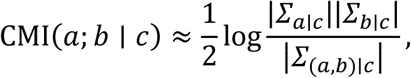

where *∑*_(·)_ denotes the conditional covariance matrices. Covariance shrinkage was applied to stabilize estimation in niches containing limited numbers of cells. HARMONIC encoded three complementary dependencies:

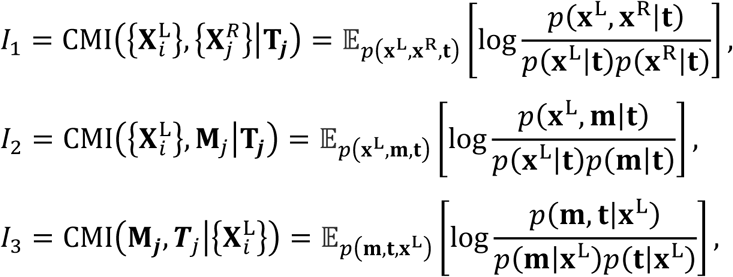

The three CMI terms capture complementary conditional associations among molecular, morphological and tissue architectural features. Because CMI does not independently indicate whether the surrounding architecture facilitates or restricts signaling, these terms are retained as separate contextual features with learnable scaling parameters. For each candidate sender-receiver cell pair, HARMONIC identifies the niches containing the two cells and incorporates the corresponding CMI features into its edge representation. Thus, cell pairs sharing the same sender and receiver niches can share niche-level contextual features while retaining distinct cell-level representations.

##### *Step3*: Integration into graph learning and CCC scoring

For each LR pair, HARMONIC applied an edge-biased graph Transformer to *G*^(L,R)^. Each cell node was initialized with a multimodal representation combining its transcriptomic profile and cellular morphology,

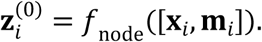

Each directed edge from sender cell *i* to receiver cell *j* was represented as

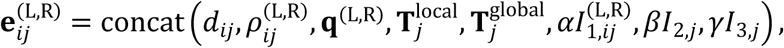

where α, *β*, and *γ* are learnable scaling parameters. This formulation exposes the model to the raw local and global architectural representations as well as the CMI-encoded dependencies, while maintaining cell-resolved molecular and morphological node states.

The edge-biased graph Transformer integrates molecular, morphological and histological context through attention-based message passing. For each candidate sender-receiver edge, an attention logit is computed from the endpoint node representations and the associated edge features. For attention head ℎ, the pre-softmax attention logit is defined as

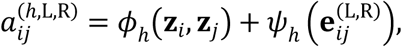

where *φ*_ℎ_ measures compatibility between the sender and receiver node states and *ψ*_ℎ_ generates a nonlinear edge bias from the LR, distance, CMI, and architectural features. For each receiver cell *j*, logits were normalized over its candidate senders *N*(*j*),

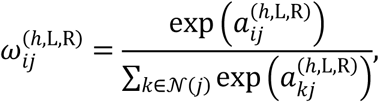

and the resulting weights were used to aggregate sender messages and update the receiver representation. Repeating this operation across layers allowed the score of a target edge to incorporate the composition and state of the surrounding local communication neighborhood.

The normalized attention weights are used to aggregate messages from candidate senders and update receiver-cell representations across Transformer layers. These weights support message passing but are not used directly as CCC scores. Instead, HARMONIC defines the LR-specific CCC score from the final-layer pre-softmax attention logits averaged across attention heads:

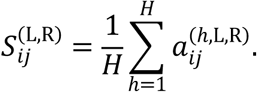

The resulting scores 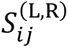 are used to rank candidate interactions for each LR pair and are linearly rescaled to [0,1] when required for visualization or downstream aggregation.

##### *Step4*: Self-supervised training strategy

HARMONIC is trained using edge-masked self-supervision with CMI-derived self-supervision. At each training epoch, a subset of candidate edges is removed from graph message passing to generate a perturbed graph. An edge decoder then predicts communication-tendency scores for the retained edges using their learned node representations, with CMI-derived scores serving as self-supervisory targets. Ranking-consistency and cross-view consistency objectives further regularize training (Supplementary section 2). After optimization, LR-specific CCC scores are obtained from the final-layer pre-softmax attention logits rather than the reconstruction decoder.

#### Output generation and downstream aggregation

After training, HARMONIC assigns an LR-specific CCC score to each directed candidate edge. Following CellNEST^6^, the top 20% of scored candidate edges are retained as high-scoring predictions for downstream analysis. CCC abundance for each LR pair is defined as the number of retained edges and can be aggregated by sender-receiver cell type, niche or signaling pathway. For three-dimensional ST, the framework is extended by including inter-slice candidate edges between spatially corresponding cells.

### 2. Simulation data creation and experimental settings

#### General settings

We generated a synthetic spatial dataset comprising 20,000 sender, receiver and noninteracting cells. Senders were positioned at the centers of circular niches, receivers in surrounding rings and noninteracting cells elsewhere. Ten ligands in sender cells and their corresponding receptors in receiver cells defined ten ground-truth LR pairs with the remaining 40 genes as non-interacting. All genes in noninteracting cells were treated as noninteracting.

For each receiver *j*, senders within 50 pixels were considered candidate interacting partners. Receptor expression was generated as

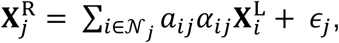

where 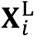 is the corresponding ligand expression, *a*_ij_ represents architecture-dependent communication, α_ij_ is the interaction weight sampled from a uniform distribution, and *ε*_j_ denotes additive Gaussian noise sampled from N(0, 0.05^2^).

Matched synthetic H&E images were generated to provide image-visible information about cellular phenotypes and tissue architecture. Sender, receiver and noninteracting cells were assigned distinct image-visible phenotypes, whereas high- and low-signal states were represented by different phenotype intensities, with low-signal cells rendered more faintly. Communication-promoting niches in scenario I and restrictive boundaries in scenario II were also explicitly represented in the images. Thus, the synthetic histological features were designed to reflect the predefined communication rules, enabling controlled evaluation of the model’s ability to incorporate architectural context rather than establishing a causal relationship between histological features and signaling in biological tissues.

#### Scenario I: recovery of underestimated communication under weak LR co-expression

In this scenario, ground-truth sender-receiver interactions were assigned either high- or low-signal states. Low-signal interactions retained their ground-truth communication status but used α ∼ U(2,4), producing a fivefold reduction in expected interaction strength relative to U(10,20). A permissiveness parameter *p* ∈ [0,1] controlled the proportion of low-signal interactions supported by the communication-promoting architecture, with the extent and density of the corresponding image-visible niche increasing with *p*. This scenario assessed whether HARMONIC could recover genuine interactions missed using molecular evidence alone.

#### Scenario II: suppression of spurious communication across architectural barriers

Here, we simulated proximal cells separated by restrictive structures, i.e., ring-shaped boundaries, which were simulated in the synthetic H&E images. A barrier-strength parameter *b* ∈ [0,1] jointly controlled boundary visibility and cross-boundary communication suppression. For sender-receiver pairs crossing a boundary, their contribution to receptor expression was scaled by 1 − *b*; thus, *b* = 0 represented no restriction and *b* = 1 complete suppression. This scenario assessed whether HARMONIC could exclude LR-supported but less anatomically plausible interactions.

### 3. PCP data collection and processing

#### Patient recruiting

19 patients (29 samples) with PCP were recruited from the Department of Neurosurgery, West China Hospital, Sichuan University, under institutional ethics approval no. 20211047A, with the informed consent obtained from all patients. PCP diagnosis was established according to the 2021 WHO classification of central nervous system tumors. Tumor specimens were obtained by surgical resection before any antitumor treatment. Formalin-fixed, paraffin-embedded tissue blocks were prepared for Xenium spatial transcriptomic profiling and CODEX imaging.

#### PCP tissue sequencing on Xenium

RNA quality of the formalin-fixed, paraffin-embedded PCP specimens was assessed using DV200, and tissue morphology was evaluated by H&E and DAPI staining before Xenium processing. Five-micrometer tissue sections were mounted on Xenium slides and processed according to the 10X Genomics Xenium user guides CG000578, CG000580, CG000582 and CG000584. Following deparaffinization and decrosslinking, sections were hybridized with the Xenium Human 5K panel at 50 °C for 20 h. Probe ligation, amplification, background quenching and DAPI staining were performed according to the manufacturer’s instructions. Imaging and transcript decoding were conducted using the Xenium Analyzer (10X Genomics; PN-1000529).

#### CODEX preparation and imaging

FFPE tissue sections adjacent to those used for Xenium profiling were analyzed by CODEX. Carrier-free monoclonal or polyclonal antibodies against human proteins were screened by multichannel immunofluorescence and conjugated to oligonucleotide barcodes using the Akoya Antibody Conjugation Kit. Additional validated conjugated antibodies were obtained directly from Akoya Biosciences. Five-micrometer sections were mounted on APTES-coated coverslips, baked at 60 °C overnight, deparaffinized in xylene and rehydrated through graded ethanol. Antigen retrieval was performed in TE buffer at pH 9 for 10 min, followed by blocking and incubation with the antibody panel for 3 h at room temperature. Multiplex imaging was performed according to the CODEX user manual using a Keyence BZ-X810 fluorescence microscope equipped with a Nikon CFI Plan Apo λ ×20/0.75 objective and the Akoya CODEX system. Raw images were stitched and processed using the CODEX Processor software.

### 4. CCC evaluation metrics

#### Contextual distance enrichment score (CDES)

To quantify the concordance of inferred CCC with a prespecified spatial and histological context prior, we defined CDES extending the original distance enrichment score^33^ (DES) to incorporate H&E-derived tissue context. For each LR pair *l* = (L, R), candidate spatially proximal cell-cell pairs were ranked using a reference score integrating LR expression, intercellular distance and histological-context similarity:

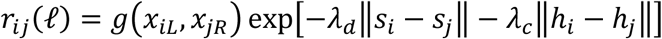

For each method, candidate edges were independently ranked by their predicted CCC scores. Agreement between the method-specific and reference rankings was quantified using a GSEA-style running-sum statistic following the original DES framework. Final CDES values were averaged across LR pairs, with higher values indicating greater concordance with the prespecified molecular, spatial and histological reference. Because histological information contributes to this reference ranking, CDES was used as a complementary context-consistency metric rather than as an independent ground-truth measure of CCC accuracy.

#### Edge-level graph stability score (EGSS)

To evaluate the robustness of CCC inference under transcriptomic or spatial perturbation, we calculated the EGSS^32^. LR pairs of each candidate cell-cell pair were ranked according to their predicted communication scores in the original and perturbed data, generating the ranked edge lists. EGSS was calculated as the Spearman rank correlation between the two complete edge lists and was linearly transformed to the interval [0,1]. Final method-level EGSS values were obtained by averaging scores across evaluated LR pairs and tissue regions. Higher EGSS values indicate greater preservation of edge-level CCC graph structure under perturbation and therefore greater inference stability.

### 5. Niche-level pathway and ligand-receptor abundance analysis

To characterize CCC programs across necrosis-associated spatial niches, niches were ordered by their distance from the nearest necrotic region. For each niche, LR - level CCC abundance was calculated from the number of retained CCC events. LR pairs were mapped to MSigDB^48^ Hallmark pathways when either the ligand or receptor gene was included in the corresponding gene set, and pathway-level CCC abundance was calculated by summing the abundance of mapped LR pairs. Migration- and proliferation-related pathways were defined based on Hallmark annotations and compared across the ordered niches. Representative LR pairs were ranked by CCC abundance within necrosis-proximal and necrosis-distal niches and visualized using dot plots.

### 6. Human ovarian cancer cell type annotation and region definition

Cell types were annotated primarily based on ST gene expression profiles using established cell-type-specific marker genes. To improve annotation accuracy in regions with high cellular heterogeneity, cellular morphology and spatial organization from the paired H&E images were additionally considered as auxiliary information. Infiltrating area was annotated on H&E as the transitional region where scattered tumor cells are admixed with non-neoplastic tissue, defined by an estimated tumor-to-normal cell ratio of ∼10-20 per 100 cells.

### 7. Cellular complexity calculation

Cellular complexity was quantified from paired H&E images using the tissue complexity metric proposed in our previous research^40^. Briefly, tissue complexity was defined as a composite score integrating two complementary histological properties: cell density and morphological heterogeneity. Cell density was estimated from the number of segmented nuclei normalized by the theoretical maximum cell count within each local image patch. Morphological heterogeneity was calculated by combining nuclear shape, size and chromatin texture variability derived from segmented nuclei. The final complexity score was computed as an equally weighted combination of the normalized density and heterogeneity scores, with higher values indicating greater cellular crowding and morphological diversity. Regions were subsequently stratified into high- and low-complexity subsets for downstream analyses.

### 8. Signaling direction analysis

To assess whether inferred CCC directions aligned with the annotated interface of tumor and immune-infiltrated area, we analyzed the orientation of sender-to-receiver signaling vectors relative to the local boundary-normal direction. For each inferred CCC event, the angle (θ) between the signaling direction vector and the boundary-normal vector directed from TAMs toward tumor cells was calculated. Directional alignment was quantified as 1 - cos θ, where lower values indicating stronger and 0° indicating perfect directional agreement. The Euclidean from each CCC event to the nearest immune-infiltrated boundary was also calculated, and directional alignment was evaluated as function of boundary distance. This analysis was used to determine whether inferred signaling directions were coordinated across the interface of tumor and immune-infiltrated area, and aligned with H&E-derived morphological gradients.

### 9. Identification of differentially expressed CCC genes

To identify genes that may be regulated by or regulate inferred CCC, we used tradeSeq^43^ to perform differential expression analysis in receiver cells. For each LR pair or signaling pathway, the received signaling abundance of each receiver cell was modeled as a continuous covariate, analogous to pseudotime-associated expression analysis. Genes were tested for expression changes with increasing received signaling abundance using the association test implemented in tradeSeq. Statistical significance was assessed using Wald tests, followed by Benjamini-Hochberg correction for multiple testing. Genes with an adjusted *P*<0.05 were considered differentially expressed genes associated with CCC. For visualization, receiver cells were ordered according to their received signaling abundance, and the scaled expression levels of selected genes were displayed as heatmaps to show their expression trends.

### 10. Anatomical region annotation in mouse kidney

Anatomical regions of the adult mouse kidney, including the cortex, OSOM, ISOM and IM, were delineated from the paired H&E image based on established renal histological features. Independently, the ST data were subjected to spatially aware clustering with K=4 to identify transcriptionally distinct spatial domains. The resulting ST-derived domains were matched to the H&E-defined anatomical regions according to their spatial distribution and characteristic regional expression patterns. The concordant region assignments were used for subsequent region-specific CCC and expression analyses.

### 11. Anatomical region-associated differential expression analysis

To provide independent transcriptional support for region-associated LR signals, we analyzed six mouse kidney ST sections from 10X Genomics. Matched H&E images were used to delineate four anatomical regions: cortex, OSOM, ISOM, and IM. ST spots were assigned to these regions according to their spatial coordinates. For each sample and anatomical region, spot-level expression values were aggregated to generate regional pseudobulk profiles. Expression values of selected region-associated genes were normalized and log-transformed for visualization, and z-scored across anatomical regions and samples for heatmap display. Regional differences in representative receptor-gene expression were assessed using Friedman tests followed by pairwise Wilcoxon signed-rank tests with Holm correction for multiple comparisons. Genes showing consistent enrichment in their expected anatomical regions across samples were used to provide independent transcriptional support for HARMONIC-inferred region-associated LR signals.

### 12. Random-walk-based downstream programs validation

To evaluate the concordance between HARMONIC-inferred signaling activity and downstream transcriptional responses, LR-specific downstream genes were derived using scSeqComm^65^. Receptor-to-transcription factor associations were obtained from TF_PPR_KEGG_human, and transcription factor-to-target relationships were obtained from TF_TG_TRRUSTv2. For each ligand-receptor pair, downstream genes were ranked according to their association with the receptor, and the top 5% of genes present in the expression matrix were retained. At the individual-cell level, downstream-gene activity was quantified as the average expression of the retained genes in each receiver cell. Pearson correlation coefficient was calculated between the HARMONIC-derived received-signal score and downstream-gene activity for each ligand-receptor pair. Spatial maps of downstream-gene activity were further compared with the corresponding CCC signals to assess spatial concordance.

### 13. TCGA ovarian cancer data processing and BayesPrism deconvolution

TCGA ovarian cancer bulk RNA sequencing data and clinical annotations were analyzed using primary tumor samples only. To obtain TAM abundance and compartment specific expression from bulk profiles, we performed deconvolution with BayesPrism^46^ using a single cell reference with curated cell type labels. BayesPrism was run with default parameters. Cell type fractions were extracted with get fraction, and predicted cell type specific expression was obtained with get exp. TAM specific expression was used for CCC signature analyses and tumor cell specific expression was used for tumor signature analyses.

### 14. Definition and scoring of TAM-tumor and tumor marker signature

The TAM-tumor signature was defined using receptor genes from the five most abundant TAM-to-tumor signaling pathways inferred by HARMONIC. For each pathway, the 20 ligand-receptor pairs with the highest CCC abundance were selected, and the receptor genes expressed in tumor cells were combined to generate the TAM-tumor signature. Duplicate genes were removed. An established tumor-cell marker signature was used as a comparator, as previously reported and listed in Table S3. For each TCGA ovarian cancer sample, signature scores were calculated using BayesPrism-inferred tumor-cell-specific expression profiles. Expression values were standardized across samples for each gene, and the signature score was calculated as the mean standardized expression of all genes within the corresponding signature. The resulting TAM-tumor CCC and tumor-cell marker signature scores were used for downstream gene-set enrichment, survival and multivariable analyses.

### 15. Stratified survival and multivariable modelling

Patients were stratified into TAM-high and TAM-low groups according to the cohort median of BayesPrism-inferred TAM abundance. Within each group, patients were further divided into signature-high and signature-low groups using the median TAM-tumor CCC signature score. Overall survival was compared between signature-high and signature-low patients using Kaplan-Meier analysis and two-sided log-rank tests. The established tumor-cell signature was evaluated using the same procedure as a comparator. To determine whether the TAM-tumor CCC signature was independently associated with survival, a multivariable Cox proportional hazards model was fitted in TAM-high patients, with adjustment for patient age and cancer grade. Associations were reported as hazard ratios with 95% confidence intervals and corresponding P values.

### 16. Gene set enrichment analysis (GSEA)

GSEA was performed using BayesPrism-inferred tumor-cell-specific expression profiles from TCGA ovarian cancer patients. Patients were divided into TAM-high and TAM-low groups according to the cohort median of TAM abundance, and genes were ranked based on differential expression between the two groups. Enrichment of the TAM-tumor CCC signature was assessed using the fgsea R package with eps = 0, minSize = 10, and scoreType = “pos”. Enrichment curves were visualized using fgsea and ggplot2.

### 17. Visualization of CCC inference results

#### Edge-based visualization

For selected LR pairs, directed CCC events were visualized by drawing arrows from sender-cell coordinates to the corresponding receiver-cell coordinates. Arrow direction represented the inferred signaling direction, and retained high-confidence CCC events were overlaid on the paired H&E image.

#### Spatial vector-field visualization

To summarize local signaling directions, cell-level sender-to-receiver CCC vectors were aggregated within a regular spatial grid. For each grid region, a representative vector was calculated from the local CCC events, with vector direction indicating the dominant signaling direction and vector magnitude reflecting the aggregated CCC score.

#### Spatial intensity-map visualization

For each receiver cell, the inferred CCC score or received-signal score was mapped to its spatial coordinate and spatially smoothed to generate a continuous intensity map. Higher intensity indicated stronger receiver-side signaling activity.

## Data availability

All publicly available datasets analyzed in this study, including Xenium and Visium HD datasets from mouse brain, mouse kidney, human glioblastoma and ovarian cancer, as well as cross-cancer transcriptomic and clinical data are summarized in Supplementary Table S6 with their source publications and download links.

The in-house papillary craniopharyngioma (PCP) datasets, including spatial transcriptomic (Xenium) data and spatial proteomic (CODEX) data, have been deposited in the repository of the National Genomics Data Center (NGDC), under accession numbers OMIX021475 and OMIX021470, respectively. These datasets can be shared for non-commercial academic research in accordance with the applicable data-access requirements.

## Code availability

Codes are available at https://github.com/JYX-111/HARMONIC.

### Acknowledgements

This work was supported by the NIHR Brain Injury MedTech Co-operative, the NIHR Cambridge Biomedical Research Centre (NIHR203312), Addenbrooke’s Charitable Trust. This work presents independent research funded by the National Institute for Health and Care Research (NIHR). The views expressed are those of the author(s) and not necessarily those of the NHS, the NIHR or the Department of Health and Social Care. This work was supported by the National Natural Science Foundation of China (32571681). Xiaofei Wang reports financial support by China Scholarship Council. Chao Li reports financial support by the Guarantors of Brain.

## Competing interests

None.

## Notes

### Competing Interest Statement

The authors have declared no competing interest.

### Summary of Updates

Update author: add Binzhi Qian, delete Yonggao Mou; add more experimental results

